# Structural Basis of CD4 Downregulation by HIV-1 Nef

**DOI:** 10.1101/2020.04.21.054007

**Authors:** Yonghwa Kwon, Robyn Kaake, Ignacia Echeverria, Marissa Suarez, Charlotte Stoneham, Peter W. Ramirez, Jacob Kress, Rajendra Singh, Andrej Sali, Nevan Krogan, John Guatelli, Xiaofei Jia

## Abstract

The HIV-1 protein Nef suppresses multiple immune surveillance mechanisms to promote viral pathogenesis^1^. Individuals infected with HIV-1 encoding defective *nef* genes do not develop AIDS for decades^2,3^. A key target of Nef is the cellular receptor CD4. Although essential for viral entry into host cells, CD4 is problematic for the virus later in its replication cycle: CD4 disrupts processing of the viral glycoprotein, Env, inhibiting infectivity^4^; it interferes with the release of new virions^5,6^; and it causes vulnerability to superinfection, causing premature cell death and limiting viral productivity^7^. Furthermore, binding of CD4 to Env exposes otherwise-concealed Env epitopes, rendering infected cells more susceptible to antibody-dependent cellular cytotoxicity and virus particles more susceptible to neutralizing antibodies^8-10^. HIV-1 has evolved strategies to mitigate these problems. Newly synthesized CD4 is targeted in the endoplasmic reticulum by the viral Vpu protein for proteasomal degradation^11^. Surface-expressed CD4, in contrast, is targeted by Nef for endocytosis and lysosomal degradation^12-15^. Nef’s effect on CD4 involves hijacking of clathrin adaptor complex 2 (AP2)-dependent endocytosis^16,17^. Although how Nef associates with a part of the tetrameric AP2 is understood^18^, a complete understanding of the interaction, especially how CD4 is sequestered by Nef into a complex with AP2, has remained elusive. Here, we present a high-resolution crystal structure that describes the underlying mechanism. An intricate combination of conformational changes occurs in both Nef and AP2 to enable CD4 binding and downregulation. Strikingly, a pocket on Nef previously identified as crucial for recruiting class I MHC is also responsible for recruiting CD4, revealing a potential approach to inhibit two of Nef’s activities and sensitize the virus to immune clearance

---

To pursue the structure, we assembled the protein complex *in vitro*. We fused the cytoplasmic domain of CD4 (CD4_CD_) to the C-terminus of Nef *via* a 36 amino acid-long, flexible linker (Fig. 1a). Instead of using the full length CD4 tail (394-433), we included only residues 394 to 419, containing all the CD4 determinants reportedly required for Nef-mediated downregulation^13,19,20^. Since Nef residues within the N-terminal amphipathic helix are dispensable for CD4 downregulation^21^, we truncated 25 amino acids from the Nef N-terminus. We engineered the tetrameric AP2 complex by removing the mobile C-terminal domain of the μ2 subunit (136-423), enabling AP2 to adopt an open conformation in which its cargo binding sites are accessible. Binding between the Nef-CD4_CD_ fusion protein and the modified AP2 construct (AP2^Δμ2-CTD^) was confirmed in a GST pulldown assay (Fig. 1b).

**Fig. 1.**
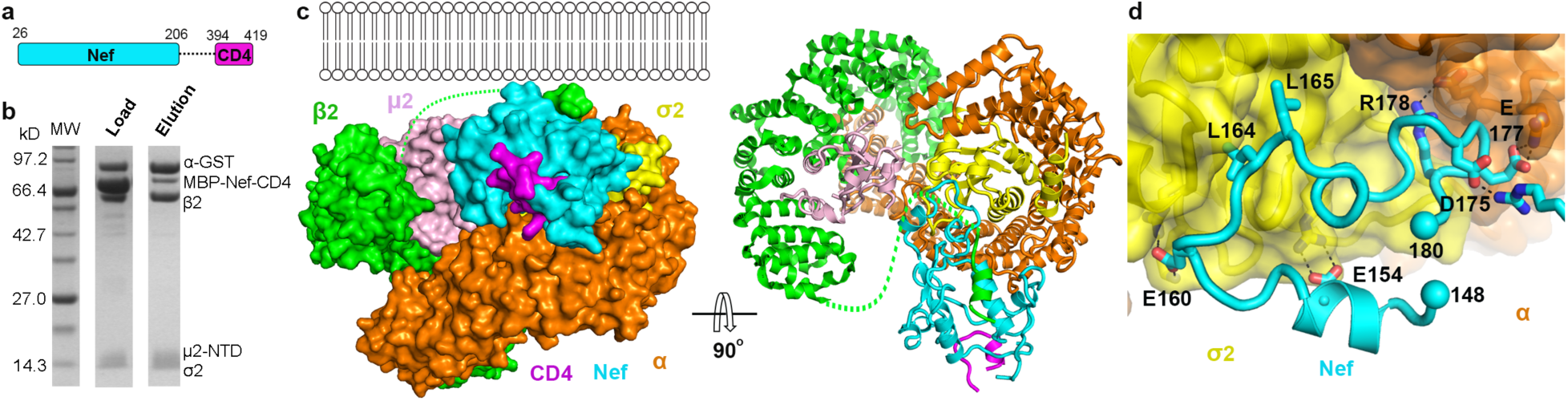
*In vitro* assembly and crystal structure of the Nef, CD4 cytoplasmic domain, and clathrin AP2 complex. **a**, Cartoon illustrating the design of the Nef-CD4_CD_ fusion protein. **b**, *In vitro* GST pulldown assay confirming the binding between the MBP-Nef-CD4_CD_ fusion protein and the AP2^Δμ2-CTD^ complex. **c**, Crystal structure of the complete protein complex in two views: along the membrane plane (left) and downward from the membrane (right). **d**, Nef associates with AP2 mainly through its C-terminal loop (148-180). The rest of Nef is not shown for an unblocked view.

We then solved the crystal structure of the Nef-CD4_CD_ fusion and AP2^Δμ2-CTD^ complex to a resolution of 3.0Å (Fig. 1c, Extended Data Table 1). All polypeptides are largely resolved except for the flexible linker between Nef and CD4_CD_, which is disordered as expected, and part of the N-terminal region of β2. As revealed by the structure, Nef functions as a “connector” between AP2 and CD4_CD_; CD4_CD_ binds Nef but none of the subunits of AP2. As previously reported, Nef’s C-terminal loop interacts with AP2 in part *via* mimicry of the acidic dileucine motifs in cellular proteins that are recognized by the α-σ2 subunits (Figure 1d)^18^. The extensive interface here, involving several charge-charge and hydrophobic interactions, is the foundation of the Nef-AP2 association.

CD4 is recruited to a pocket on Nef that is opposite the C-terminal loop. The association is mainly hydrophobic and involves three CD4 residues: Ile410, Leu413, and Leu414 (Fig. 2a). The dileucine motif of CD4 – Leu413/414 – is within a short helix. Leu414 and Ile410 dip into a hydrophobic pocket of Nef formed by Phe121, Trp124, Met79, Thr138, and Pro78 (Fig. 2b). Leu413 of CD4 sits just outside of that pocket and is accommodated by Nef residues Phe121, Leu37, Asn52, and Cys55 (Fig. 2b). Nef residue Asp123 contributes to CD4 binding by hydrogen-bonding with the backbone nitrogen of CD4 Leu413 and supports the helix-turn of CD4 (Fig. 2b). These structural findings explain previously observed roles of CD4 residues – the dileucine motif and Ile410^13,20,22^ – as well as Nef residues F121, D123, and Trp124^23,24^. To further confirm the structural model, we tested the mutations F121D, D123R, M79D, and T138D for their effects on CD4 surface-downregulation. The Nef mutants F121D and D123R were expressed similarly to the wild type but were unable to downregulate CD4 (Fig. 2ef). The M79D and T138D Nef mutants were relatively poorly expressed, but dose-response experiments suggested that this was insufficient to explain their functional defects (Fig. 2ef and Extended Data Fig. 1).

**Fig. 2.**
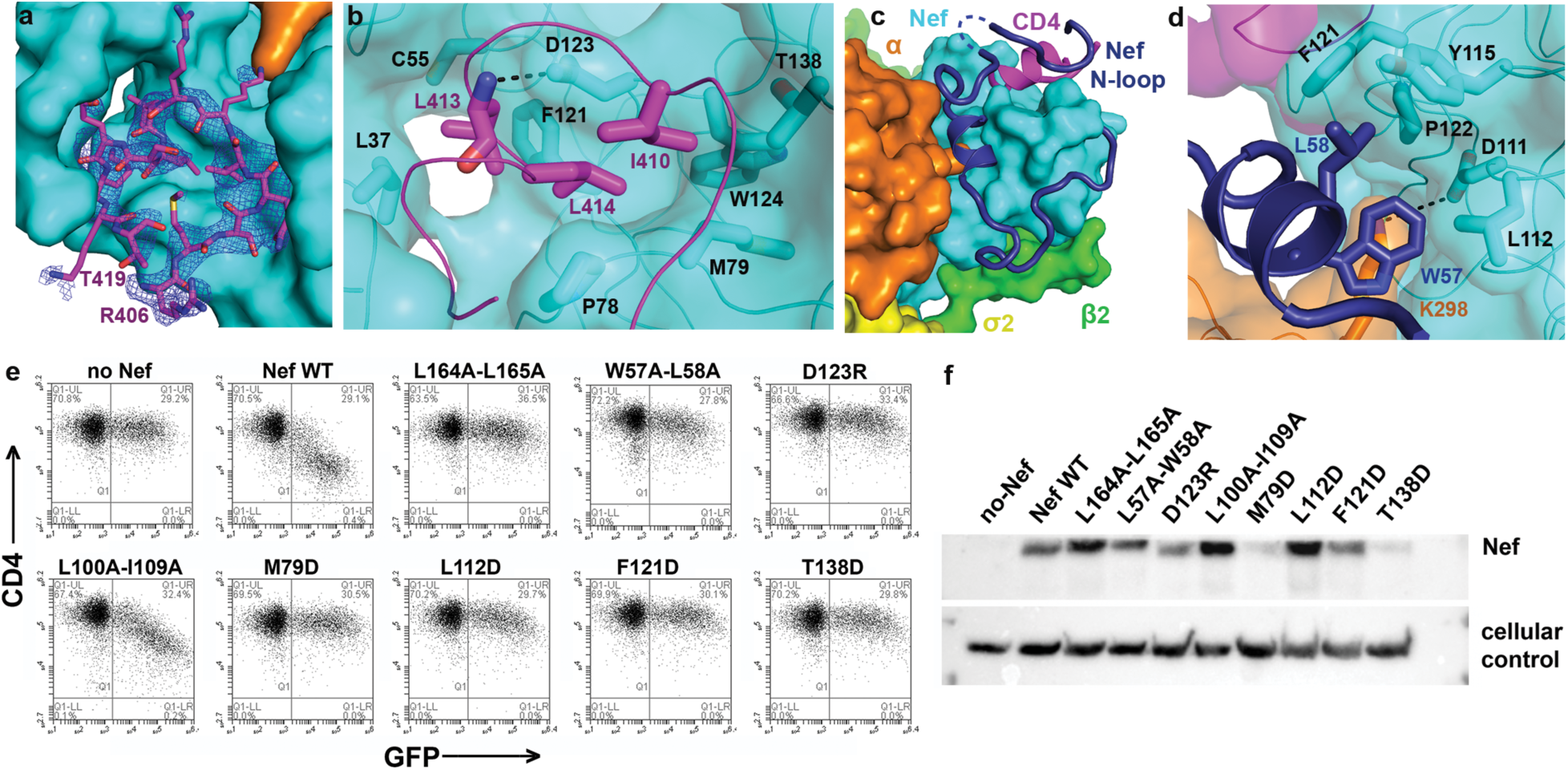
Recruitment of CD4 cytoplasmic domain and the role of Nef N-terminal loop. **a**, a short stretch of the CD4_CD_ binds to a pocket on Nef. Electron density of the CD4_CD_ (blue mesh, 2Fo-Fc map at 1.0s with B factor sharpening of −50Å^2^) is shown. **b**, Hydrophobic interactions mediated by CD4 I410, L413, and L414. **c**, Nef N-terminal loop (dark blue) adopts a unique conformation in the complex. Part of the loop forms a wall of the CD4-binding pocket. Dotted line represents Nef 41-46 that are disordered in the structure. **d**, Nef residues W57 and L58 mediate the docking of the short helix into the hydrophobic pocket formed at the Nef-α interface. **e**, CD4 downregulation by Nef mutants was measured using surface-staining and flow cytometry; GFP is a transfection marker. **f**, Expression of Nef mutants in the transfected cells used in the CD4 downregulation assay measured by western blot; the cellular control is β-actin.

CD4 recruitment is secured by the Nef N-terminal loop, which is highly ordered in the structure (Fig. 2c and Extended Data Fig. 2). The part of the loop immediately connected to the rigid core of Nef, residues Phe68 to Pro75, wraps around the core. N-terminally, the loop then takes a sharp “U-turn”, placing a short helix at the interface between the Nef core and the α subunit of AP2. Two residues within this helix, Trp57 and Leu58, fit into a hydrophobic pocket formed mainly by Nef residues: Leu112, Pro122, Phe121, and Tyr115 (Fig. 2d). Lys298 of α, with its head group stabilized by Nef Asp111 through charge-charge interactions, also contacts Nef Trp57 *via* its hydrocarbon chain (Fig. 2d). The Nef acidic cluster (Glu62-65), located in the “U-turn” region, does not contribute to CD4-binding, consistent with previous functional observations^16,25,26^. The Nef N-terminal loop extends to contact the CD4 tail (Fig. 2c). This part of Nef, residues Cys55 to Val33, forms a wall-like structure to support CD4-binding. Here, contacts are made by several Nef residues with the short helical turn of CD4 and flanking residues. Overall, this conformation of the Nef N-terminal loop explains the crucial roles of Trp57, Leu58, Leu112, Tyr115, and Pro122 in CD4-downregulation^16,17,21,23,27^ (Fig. 2ef) as well as the cooperativity in the three-way binding between CD4_CD_, Nef and the α/σ2 hemicomplex of AP2^28^. Nef residues 41-46 are disordered with most of their electron density missing (Fig. 2c, dotted line). The density used for modeling Nef 34-40 is significant, but isolated and less well-defined, yielding a degree of uncertainty in the structural assignment for this region (Extended Data Fig. 2). We challenged the assignment by mutating Leu37 to Asp, as the model predicts that Leu37 directly contacts CD4 Leu413 (Fig. 2b). This mutation abolished CD4 downregulation (Extended Data Fig. 1), supporting the structural assignment and the role of Leu37 in CD4 downregulation.

Nef binding causes a large conformational change and severe destabilization in the β2-μ2 half of AP2. Overlaying the α and σ2 subunits of the “open” AP2 structure^29^ on the current structure reveals that the β2 and μ2 N-terminal domains (μ2-NTD) of AP2 in the current structure “move” out as one rigid body, causing a greater “opening” of the tetramer (Fig. 3a). In addition, a large portion of the β2 N-terminus, residues 1-88 encompassing the first four helices, is displaced from the rest of β2. If β2 had maintained its original fold, then its N-terminus would clash with the Nef core (Extended Data Fig. 3); the observed displacement is likely necessary to accommodate Nef-binding. While most of this displaced portion of β2 becomes disordered, the first helix preserves its helical structure and binds to the Nef core through hydrophobic interactions (Fig. 3bc).

**Fig. 3.**
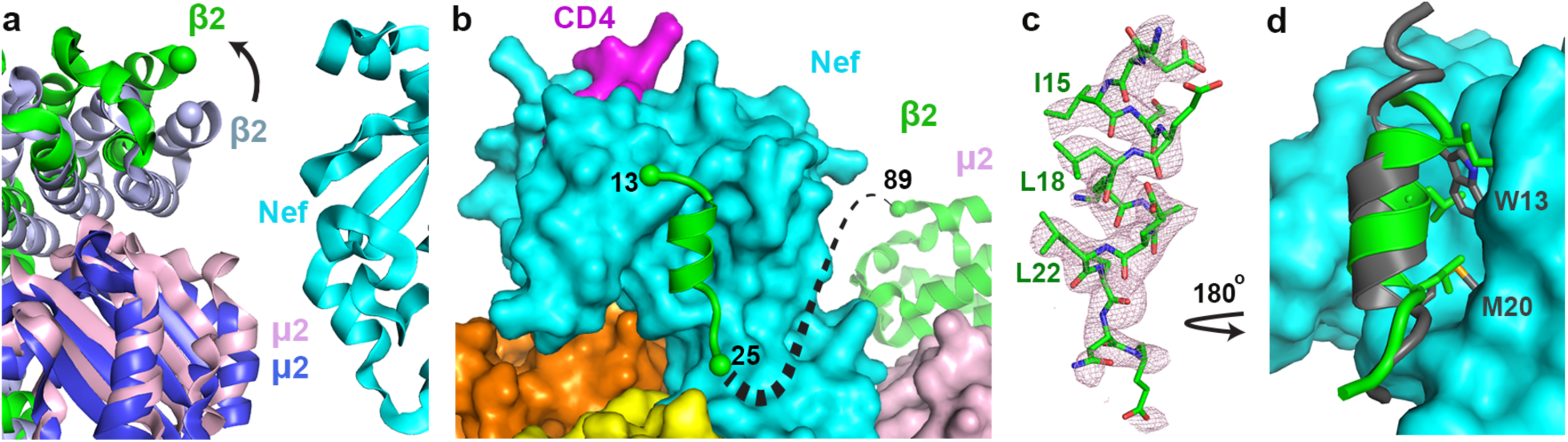
Nef binding induces conformational change in the β2 subunit of AP2. **a**, Current structure is overlaid, on the α-σ2 half, with the open AP2 (PDB ID: 2XA7). β2 (green) and μ2 (pink) subunits in the current structure move outward in comparison to β2 (light blue) and μ2 (blue) subunits of the unbound AP2. **b**, The first helix of β2 binds to the Nef core, while the next three helices become disordered. **c**, Density for the first helix of β2 (pink mesh, 2Fo-Fc map at 1.0s with B factor sharpening of −50Å^2^) is shown. Residues important for association with Nef core are labeled. **d**, Overlay of Nef in the current structure and that when in complex with MHC-I cytoplasmic domain and μ1-CTD of AP1 (PDB ID: 4EMZ) shows that the Nef N-terminal helix (gray) and the first helix of β2 (green, current structure) occupy the same site on the Nef core. Trp13 and Met20 residues in the Nef N-terminal helix and important for the intramolecular helix-core association are shown.

The Nef-induced destabilization and structural changes of the β2-μ2 half of AP2 were further characterized by chemical cross-linking mass spectrometry (XL-MS) and integrative structure modeling (Extended Data Fig. 4). Here, disuccinimidyl sulfoxide (DSSO), a MS-cleavable, bifunctional amine-reactive small molecule, was used to cross-link proximal Lys residues or N-termini of the Nef-CD4_CD_ fusion and AP2^Δμ2-CTD^ complex. Cross-linked proteins separated by SDS-PAGE were trypsin digested and resulting peptides analyzed by specialized LC-MS^3^ experiments for identification of cross-linked residues (Extended Data Fig. 4a). Application of this pipeline to the complex identified intra- and inter-linked peptides corresponding to 90 unique cross-linked residues (Extended Data Table 2, Extended Data Fig. 4b). Lys residues from the displaced β2 N-terminal domain (1-88) are involved in a total of 24 cross-links (Extended Data Table 2, bold entries). Importantly, most of these cross-links are made by Lys residues flanking the first helix (i.e. Lys5, 11, 12, 26, and 27), consistent with this helix binding specifically in the complex. In contrast, Lys residues from the other dislocated β2 N-terminal helices (i.e. Lys29, 31, 35, 36, 45, 66, 67, 78) are rarely observed, consistent with a lack of fixed residence within the complex. The XL-MS data, which captures structural information from a conformationally heterogeneous population of protein complexes as they exist in solution, the crystal structure, and other structural information (*methods*) were used for integrative modeling^30^ to produce a model ensemble which describes the complex in full (Extended Data Fig. 4c). Segments that are disordered in the crystal structure are modeled and represented as helices or flexible strings of beads (Extended Data Fig. 4c). The model ensemble agrees satisfactorily with the observed cross-links (89% of cross-links are satisfied, Extended Data Fig. 4d). Furthermore, it indicates large variability in the positions and orientations of the helices in the partially unfolded β2 segment (Extended Data Fig. 4e-g), consistent with the structural heterogeneity of this region indicated by the crystallographic data.

Binding of the first helix of β2 to the Nef core in the current structure closely resembles how the N-terminal amphipathic helix of Nef (deleted from the construct used herein) binds the same location when Nef hijacks AP1 for MHC-I downregulation (Fig. 3d)^31^. That interaction, mediated by Nef Trp13 and Met20, is critical for MHC-I downregulation but is dispensable for CD4 downregulation^21,31,32^. Our current structure indicates that intramolecular association of this Nef helix with the Nef core would force the Nef N-terminal loop to deviate from the conformation associated with CD4-binding (Extended Data Fig. 5). We suspect that this is an intricate plot of the virus: Nef binding forces the N-terminal helices of β2 to destabilize and unfold. By providing a binding-site for the first β2 helix on its core, Nef provides partial compensation for this destabilization. Moreover, the N-terminal β2 helix is now used to compete the Nef amphipathic helix off the core, freeing the N-terminal loop of Nef to adopt the conformation shown in Figure 2c for CD4 downregulation. A potential caveat to this hypothesis is that mutations of Nef residues F90, L97, L100, I109, W113, and I114, which line the binding site on the Nef-core for the β2 helix, have minimal effects on the downregulation of CD4 (Fig. 2, Extended Data Fig. 1 and 6). However, since these mutations likely affect the binding of the β2 helix and the Nef N-terminal amphipathic helix similarly, they might have minimal influence on the competition between these two helices for the Nef core *in vivo*, consistent with their lack of effects on CD4 downregulation. In contrast, and consistent with the model above, mutations of this site affect the downregulation of class I MHC (Extended Data Fig. 7); alanine substitution of L100 together with I109 is particularly informative: this mutation yields a well-expressed Nef that cannot downregulate class I MHC yet downregulates CD4.

Comparing the current structure with our earlier structure of Nef in complex with the μ1-CTD and the MHC-I cytoplasmic domain reveals versatility and specificity in how the structurally homologous AP1 and AP2 are selectively co-opted to downregulate MHC-I and CD4 (Fig. 4a). For MHC-I downregulation, Nef interacts solely with the μ1 subunit and exploits the conserved Tyr-based motif-binding site on AP1^31^. For CD4 downregulation, Nef exploits the acidic dileucine-binding site on the *α* and σ2 subunits of AP2 and contacts all subunits except μ2. Our models show that by allowing the N-terminal Nef helix and the N-terminal β2 helix to bind the same pocket on the core, Nef creates a “molecular switch” that links its use of different AP complexes with the modulation of different targets: binding to AP2 frees the N-terminus of Nef from the core to recruit CD4, whereas binding to AP1 leaves the core free to bind the N-terminus of Nef and thereby facilitates the recruitment of MHC-I.

**Fig. 4.**
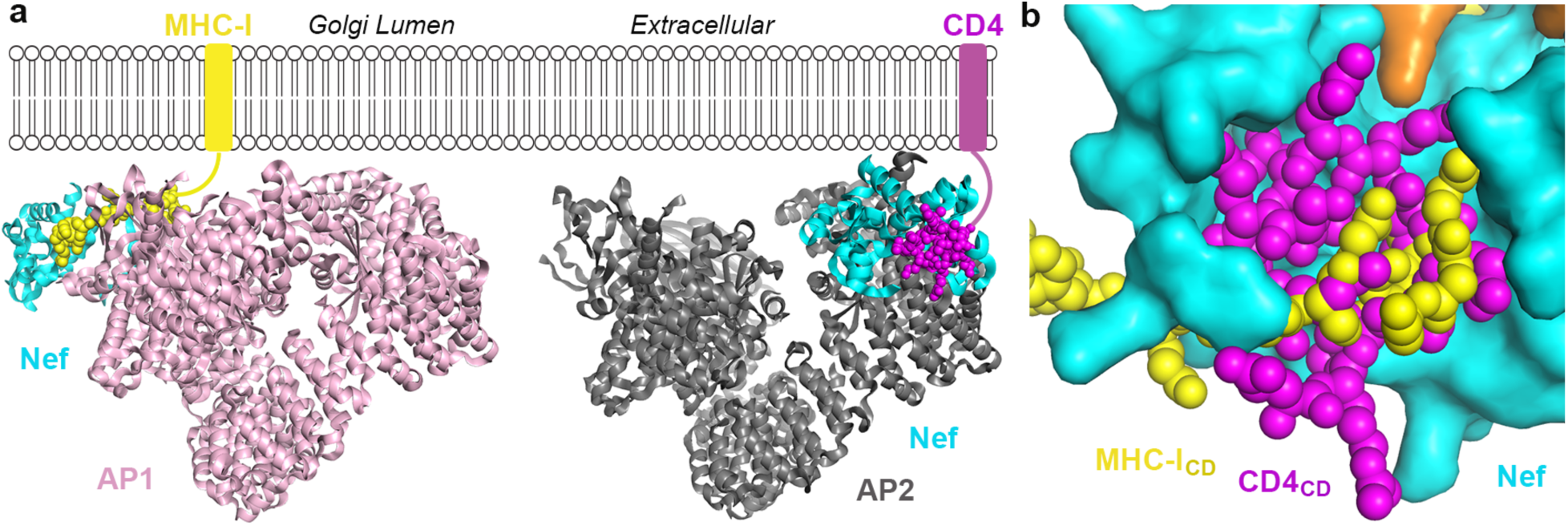
The downregulation of MHC-I and CD4 by Nef are distinctive both mechanistically and structurally, yet their cytoplasmic domains share a common binding site on Nef. **a**, Structural comparison of Nef/AP1/MHC-I_CD_ (PDB ID:4EMZ; other AP1 subunits modeled in based on overlay of the μ1-CTD) and Nef/AP2/CD4_CD_ (current structure) shows that Nef uses different ways to hijack AP1 (left) and AP2 (right) for downregulating MHC-I and CD4, respectively. **b**, Two structures, Nef/μ1/MHC-I_CD_ (PDB ID: 4EMZ) and Nef/AP2/CD4_CD_ (current structure), are overlaid on Nef (cyan). Binding of CD4_CD_ (magenta) and binding of MHC-I_CD_ (yellow) involve the same pocket on Nef.

Despite these distinct modes of binding, the cytoplasmic domains of MHC-I and CD4 share much of the same binding “pocket” on Nef (Fig. 4b). Small molecules that bind this conserved site (Extended Data Fig. 8) might block both MHC-I and CD4 surface downregulation, potentially revitalizing immune mechanisms to combat and even clear HIV-1.

## METHODS

### Materials

Gene of rat α adaptin is a kind gift from Dr. Juan Bonifacino (NIH). Genes of β2, μ2, and σ2 adaptins were amplified from a cDNA library of human HEK293T cells. For cross-linking mass spectrometry analysis, anhydrous dimethyl sulfoxide (DMSO), disuccinimidyl sulfoxide (DSSO), MS-grade trypsin, HPLC-grade water, formic acid, and acetonitrile were all purchased from Thermo Fisher Scientific. 4-20% TGX SDS-PAGE gels were purchased from Bio-Rad. MS-safe AcquaStain was purchased from Bulldog Bio.

### Cloning, expression, and purification of proteins

The Nef-CD4_CD_ fusion was constructed by fusing CD4 (394-419) to the C-terminus of HIV-1 Nef (26-206, NL4.3) *via* a flexible linker of 36 amino acids (GVDGSDEASELACPTPKEDGLAQQQTQLNLRGSGSG). The encoding gene was cloned into a pMAT9 expression vector. The fusion protein was over-expressed in *E. coli* NiCo21(DE3) cells, carrying a N-terminal maltose binding protein (MBP) tag. Cells were induced with 0.1 mM isopropyl *β*-d-thiogalactopyranoside (IPTG) at OD600 of 0.8 and grown at 16°C overnight. Cells were lysed using sonication. Expressed protein was first purified using the MBP affinity column. For GST pull-down experiments, the MBP-tagged protein was further purified by a HiTrap Q anion exchange column, followed by a final Superdex 200 size exclusion column. For crystallization, the MBP fusion tag was cleaved off by the SARS-CoV M^pro^ protease^33,34^ after the affinity purification. The tagless protein was similarly purified using HiTrap Q anioin exchange column and the Superdex 200 size exclusion column.

For AP2^Δμ2-CTD^, genes for each of the four subunits were cloned into two duet vectors: a pETDuet vector with human β2(1-591) and human μ2(1-135) at each of the multiple cloning sites, respectively; a pCDFDuet vector with rat α(1-621) carrying a C-terminal GST tag and human σ2(1-142). The heterotetrameric AP2^Δμ2-CTD^ core was expressed overnight at 22°C in the NiCo21(DE3) cells in Terrific broth after induction with IPTG. For binding assays, the GST-tagged the AP2^Δμ2-CTD^ complex was purified by a sequence of Ni-NTA gravity column, GST affinity column, and Superdex 200 size exclusion column. For crystallization, the GST tag was cleaved off by the Tev protease after the GST affinity purification, followed by a final purification using the Superdex 200 size exclusion column.

### *In vitro* GST pulldown assay

Purified proteins AP2^Δμ2-CTD-^GST (0.2 mg) and MBP-Nef-CD4_CD_ (0.4 mg) were mixed in a final volume of 100 ul and incubated at 4°C for 30 minutes. The protein solution was then loaded onto a small gravity flow column containing 0.2 ml GST resin. Flow through was collected and the resin was extensively washed with 5 × 0.9 ml GST binding buffer (50 mM Tris, pH 8, 100 mM NaCl, 0.1 mM TCEP). The bound proteins were then eluted with 5 × 0.1 ml GST elution buffer containing 10 mM reduced glutathione. The eluted proteins were analyzed by SDS-PAGE.

### Crystallization and crystallographic data collection

Crystallization was carried out using the microbatch under-oil method. The purified AP2^Δμ2-CTD^ core and the Nef-CD4_CD_ fusion chimera were mixed at 1:5 molar ratio to a final concentration of 2.5 mg/ml (25 mM Tris, pH 8.0, 100 mM NaCl, 0.1 mM TCEP, 0.1 mM PMSF). Equal volumes of the protein solution and the precipitant solution (100 mM HEPES, pH 6.9, 200 mM KCl, 15% PEG4000, 6% 1,6-hexanediol) were mixed. The drop was sealed using a mixture of paraffin and silicon oil at a 2:1 ratio.

Crystals appeared within 24 h at room temperature and grew to full size in about a week. Crystals were cryo-protected using the precipitant solution containing 20% glycerol and then frozen in liquid nitrogen. Datasets were collected at NE-CAT (24-ID) at the Advanced Photon Source, Argonne National Laboratory, and FMX/AMX (17-ID) at the National Synchrotron Light Source II, Brookhaven National Laboratory. Diffraction data was processed using HKL2000^35^. The crystals were in the P41 space group and diffracted to a highest resolution of 3.0 Å. The statistics are summarized in Table S1.

### Structure determination and refinement

The structural solution was obtained by molecular replacement using *PHASER*^36^ in *PHENIX*^*37*^. Only one molecule exists in the asymmetric unit. The PDB of the open AP2 core (2XA7) was divided into two search models: the α and σ2 hemicomplex and the β2 and μ2-NTD hemicomplex. Together with the Nef structure (4EMZ), the three models were used sequentially to successfully obtain the solution. Iterative rounds of model building in *COOT*^38^ and refinement in *Phenix*^*39*^ were carried out. The final model has an *R*_*work*_/*R*_*free*_ of 0.241/0.277. A Ramachandran plot showed that 96.4% of the residues are in the favored region, together with 3.5% in the allowed region and 0.1% as outliers. The refinement statistics are summarized in Extended Data Table 1.

### CD4 downregulation assays

HeLa cells expressing CD4 (TZM-bl, obtained from Dr. John Kappes *via* the NIH AIDS Reagent Program) were transfected using Lipofectamine2000 (Thermo Fisher Scientific) with pCG-GFP (a gift from Dr. Jacek Skowronski) and pCI-NL, a pCI-neo-based plasmid (Promega) expressing Nef_NL4-3_ or the indicated Nef-mutants. Unless otherwise indicated, 1.6 µg of total plasmid DNA was used in each transfection, 0.4 µg of pCG-GFP and 1.2 µg of pCI-NL or derivative-mutants. In some experiments, the amount of pCI-NL plasmid encoding wild type Nef was reduced in a dose-response format to allow comparison with relatively poorly expressed Nef-mutants; in those cases, the total amount of plasmid DNA was equalized with the empty vector pCI-neo. One day later, half the cells were stained for surface CD4 (anti-human CD4, Biolegend, conjugated directly to APC), fixed in formaldehyde, then analyzed by two-color flow cytometry using an Accuri 6 cytometer. “Live cell” gates were set using untransfected cells; gates for GFP were set using cells transfected only with pCI-neo; and gates for CD4 were set using cells stained with an APC-conjugated antibody isotype control. The other half of the cells were lysed in Laemmli buffer, and the proteins were resolved on 10% denaturing SDS-PAGE gels before transfer to polyvinylidene difluoride membranes. Nef was detected using a polyclonal antiserum raised to NL4-3 Nef in sheep (a gift from Dr. Celsa Spina, University of California San Diego). β-actin was detected using a murine monoclonal antibody (Sigma-Aldrich). Species-specific secondary antibodies conjugated to HRP were visualized using Western Clarity detection reagent (Bio-Rad). Chemiluminescence was recorded using a ChemiDoc Imager System (Bio-Rad).

### Class I MHC downregulation assays

HEK293 cells, which naturally express HLA-A2, were transfected using Lipofectamine2000 with pCG-GFP (0.4 µg) and pCI-NL, the indicated Nef-mutants, or the empty vector pCIneo (1.2 µg). One day later, half the cells were stained for surface HLA-A2 (Biolegend, anti-HLA-A2 conjugated directly to APC), fixed in formaldehyde, then analyzed by two-color flow cytometry using an Accuri 6 cytometer. “Live cell” gates were set using untransfected cells; gates for GFP were set using cells transfected only with pCI-neo; and gates for HLAA2 were set using cells stained with an APC-conjugated antibody isotype control. The other half of the cells were processed for western blot as above.

### DSSO-based cross-linking mass spectrometry analysis (XL-MS)

Individual preparations of Nef-CD4_CD_-AP2^Δμ2-CTD^ complex (2.8mg/mL and 0.7mg/mL at 1:5 Nef-CD4_CD_ molar excess) were cross-linked using increasing molar ratios of DSSO (Thermo Fisher Scientific), for 5, 10, or 30 minutes at 4 or 37°C. Cross-linked proteins were separated on 4-20% TGX gradient SDS-PAGE gels (Bio-Rad), stained with MS-safe AcquaStain (Bulldog Bio), and cross-linked product bands excised and submitted for in gel reduction, alkylation, and trypsin digestion. Extracted peptides were separated online by Thermo Easy nLC 1000 by reverse-phase HPLC (75 μm × 30 cm fused silica packed with 1.9-μm Reprosil-Pur C18 AQ resin (Dr. Maisch-GmbH) column), running a linear gradient of 5-30% B in 50min, 35-95% B in 5 min, and 95% B for 4 min at a flow rate of 300 nL/min (buffer A: 100% H_2_O/0.1% FA; buffer B: 100% CAN/0.1% FA). For each sample, XL-MS^3^ data was acquired on Thermo Orbitrap Elite using two similar data dependent acquisition experiments^40^ where a single acquisition cycle consisted of: 1) one full MS^1^ scan (350–1500 m/z, 120,0000 resolution, AGC target of 1×10^6^); 2) top two data-dependent MS^2^ scans (15,000 resolution, AGC target of 5×10^4^, normalized collision energy = 22%); and 3) top three (or four) MS^3^ scans (ion count target 10^4^, normalized collision energy = 35%).

Precursor ions (charge state ≥4+) were dynamically excluded for 20 seconds (tolerance of 10 ppm). Charge state and dynamic exclusion were applied to MS^2^ but turned off for MS^3^ acquisition.

Raw data was extracted to MGF format using MSConvert^41^, with MS^3^ data used for protein and peptide searches. Searches were performed by batch-tag feature of a locally installed version of Protein Prospector (v. 5. 19. 1, University of California San Francisco), with DSSO remnant mass modifications set as variable modifications (e.g. Alkene, Sulfenic-acid, and Thiol) ^40^. Peptide reports were generated using the Search Compare feature of Protein Prospector, and dead-end, intra-linked, and inter-linked peptides identified by in house software program XL-Discoverer (part of new XLTools suite)^42^. Summarization and confidence assignment of inter-linked peptides was performed by in house scripts that reduce ambiguous assignments and distribute redundant counts.

### Integrative structure modeling of the Nef-CD4_CD_-AP2^Δμ2-CTD^ complex

We applied an integrative structural modeling approach^30,43-45^ to characterize the structure of the Nef-CD4_CD_-AP2^Δμ2-CTD^ complex in solution, based on the crystal structure and the 90 DSSO cross-links. Integrative structure determination proceeded through the standard four stages^43,44,46-49^: 1) gathering data, 2) representing subunits and translating data into spatial restraints, 3) configurational sampling to produce an ensemble of structures that satisfies the restraints, and 4) analyzing and validating the ensemble structures and data. The integrative structure modeling protocol (*ie*, stages 2, 3, and 4) was scripted using the *Python Modeling Interface* (PMI) package, a library for modeling macromolecular complexes based on our open-source *Integrative Modeling Platform* (IMP) package^44^, version 2.8 (https://integrativemodeling.org). Files containing the input data, scripts, and output results are available at https://github.com/salilab/Nef_CD4_AP2.

<u>(1) Gathering data:</u>

Modeling was based on the crystal structure, a comparative model of the β2 subunit 24-89 region built based on the AP2 structure^29^ using MODELLER^50,51^ and the 90 DSSO cross-links.

<u>(2) Representing subunits and translating data into spatial restraints:</u>

To maximize computational efficiency while avoiding using too coarse a representation, we represented the Nef-CD4_CD_-AP2^Δμ2-CTD^ complex using a coarse-grained one residue per bead representation. The regions absent from the crystallographic structure and the comparative model were represented by a flexible string of beads corresponding to one residue each. To explore the positions and orientations of the Nef-CD4_CD_-AP2^Δμ2-CTD^ components, we defined the following rigid bodies: α-σ2, β2(89-591)-μ2, Nef-CD4_CD_-β2(15-23), and the 4 helices in the partially unfolded β2 segment (29-43, 49-61, 64-78, and 81-86). With this representation in hand, we next translated the input information into spatial restraints as follows.

First, the 90 DSSO cross-links were used to construct a Bayesian term that restrained the distances spanned by the cross-linked residues^52,53^. The cross-link restraints were applied to the one residue-per-bead representation for the X-ray structure, comparative models, and flexible strings of beads. Second, to use the crystal structure of the hexamer as a template, we imposed “structural equivalence” distance restraints between pairs of residues closer than 8.0 Å across an interface between two rigid bodies, designed to restrain the model to resemble the template as much as possible. Third, excluded volume restraints were applied to all pairs of beads^46,54^. Fourth, we also applied the sequence connectivity restraint, using a harmonic upper bound on the distance between two consecutive beads in a subunit, with a threshold distance equal to four times the sum of the radii of the two connected beads^46,54^.

<u>(3) Configurational sampling to produce an ensemble of structures that satisfies the restraints:</u>

The initial positions and orientations of rigid bodies and positions of the beads in the flexible strings of beads were randomized. The structural models were generated using Replica Exchange Gibbs sampling, based on the Metropolis Monte Carlo (MC) algorithm^53,55^. Each MC step consisted of a series of random transformations (*ie*, rotations and translations) of the positions of the beads and rigid bodies. The sampling produced 4,000,000 models from 80 independent runs.

<u>(4) Analyzing and validating the ensemble structures and data:</u>

Model validation follows four major steps^30,56^:

i. Selection of the models for validation: The ensemble of models for further analysis was objectively defined as follows. For each trajectory, the MC step at which all data likelihoods and priors are equilibrated (run equilibration step) was computed and all prior frames were discarded^57^. Sampling of the Nef-CD4_CD_-AP2^Δμ2-CTD^ complex yielded 2,007,800 representative structures that sufficiently satisfied the input restraints.
ii. Estimation of sampling precision: The precision at which sampling sampled the selected structures (sampling precision) was estimated^56^. The sampling precision must be comparable or higher than the precision of the structure ensemble consistent with the input data (model precision). As a proxy for testing the thoroughness of sampling, we performed four sampling convergence tests as described^56^. We performed three sampling convergence tests: 1) verify that the scores of refined structures do not continue to improve as more structures are computed. 2) Confirm that the selected structures in independent sets of sampling runs (Sample A and Sample B) satisfy the data equally well. The non-parametric Kolmogorov-Smirnov two-sample test (two sided) indicates that the difference between the two score distributions is insignificant (p-value (0.08)>0.05). In addition, the magnitude of the difference is small, as demonstrated by the Kolmogorov-Smirnov two-sample test statistic (D=0.02). 3) Cluster the structural models and determine the sampling precision at which the structural features can be interpreted. Three criteria were used for determining the sampling precision, evaluated as a function of the RMSD clustering threshold. First, the p-value is computed using the *χ*2-test (one*-*sided) for homogeneity of proportions. Second, an effect size for the *χ*2-test is quantified by the Cramer’s *V* value. Third, the population of structures in sufficiently large clusters (containing at least ten structures from each sample). Clustering is done at the RMSD threshold at which three conditions are satisfied (*χ*2-test p-value (1.0) > 0.05, Cramer’s *V* (0.0) < 0.10, and the population of clustered structures (0.96) > 0.80). The output of this protocol is a single distinct cluster containing the majority (96%) of the individual models. The sampling precision is defined as the average bead RMSD between the models within the cluster and its corresponding centroid in the finest clustering for which each sample contributes models proportionally to its size (considering both significance and magnitude of the difference) and for which a sufficient proportion of all models occur in sufficiently large clusters. The sampling precision for our integrative model is 13.5Å.
iii. Estimation of model precision: The most explicit description of model uncertainty is provided by the set of all models that are sufficiently consistent with the input information (*i*.*e*. the ensemble). For example, if the models in the ensemble are clustered into a single cluster as in the case of the Nef-CD4_CD_-AP2^Δμ2-CTD^ ensemble, the model precision is defined as the RMSD between models in the cluster. The precision for the Nef-CD4_CD_-AP2^Δμ2-CTD^ model is 8.3 Å.
iv. Quantification of the degree to which a model satisfies the information used to compute it: An accurate structure needs to satisfy the input information used to compute it. A DSSO cross-link restraint is satisfied by a cluster of models if the corresponding Cα–Cα distance in any of the models is less than 30 Å^58^. The ensemble satisfies 89% of the XLs used to compute it, including all of the Nef XLs (Extended Data Fig. 4). The unsatisfied crosslinks mostly span residues between CD4_CD_ and the NTD’s of the α and β2 subunits (Extended Data Table 2). These violations can be rationalized as non-specific interactions between CD4_CD_ and the AP2 complex when CD4_CD_ is not bound to Nef in solution, false-positive crosslinks, sample heterogeneity, insufficient conformational sampling, and coarse-grained representation of the modeled components. The remaining restraints, including structural equivalence, excluded volume, and sequence connectivity restraints, are also satisfied within their uncertainties.

To compare the X-ray structure to the ensemble obtained using integrative modeling, we computed the distribution of the Cα root-mean-square deviation (RMSD) between the X-ray structure and each of the models in the ensemble. The mean Cα RMSD is 7.1 Å (4.3-12.2 95% CI).

To indicate the most flexible parts of the structure, we assessed the uncertainty of the position and orientation of each rigid body representing the hexamer in the model ensemble. To this end, all models were superimposed on each rigid body in turn, followed by computing the average RMSD for each of the other rigid bodies (Extended Data Fig. 4e). The model ensemble indicates large variability in the positions and orientations of the 4 helices in the partially unfolded β2 segment (Extended Data Fig. 4eg), consistent with the structural heterogeneity of this region indicated by the relative lack of electron density from crystallography.

## Data availability

The coordinates and structural factors for the crystal structure have been deposited at the Protein Data Bank (PDB) with the accession code 6URI.

## Acknowledgments

We thank Yong Xiong for helpful discussions and valuable inputs. We thank the beamline staff at the Advanced Photon Source beamline 24-ID and the National Synchrotron Light Source beamline 17-ID. We thank Juan Bonifacino for providing the gene of rat α adaptin. This work was supported by the University of Massachusetts Dartmouth startup fund (X.J.), US National Institutes of Health (NIH) grants AI102778 and AI129706 (J.G.). A.S. was supported by NIH grants U19AI135990, R01GM083960, P41GM109824, and S10OD021596. N.K. was supported by NIH grants P50GM082250 and U19AI135990. R.K. was supported by NIH fellowship F32AI127291.

## Author contributions

Y.K. performed protein expression, purification, binding assay, and crystallization. Y.K. and X.J. performed data collection, structure determination, model building, and refinement. J.K. and R.S. contributed to protein expression and purification. M.S., C.S., and P.R. performed CD4 and MHC-I downregulation assays and mutagenesis. R.K. performed crosslinking-MS. I.E. performed integrative modeling. Y.K., R.K., I.E., J.G., and X.J. designed the experiments. All contributed to data analysis. J.G. and X.J. supervised the project. Y.K., J.G., and X.J. wrote the manuscript.

## Correspondence and requests for materials

should be addressed to X.J.

---

**Extended Data Fig. 1.**
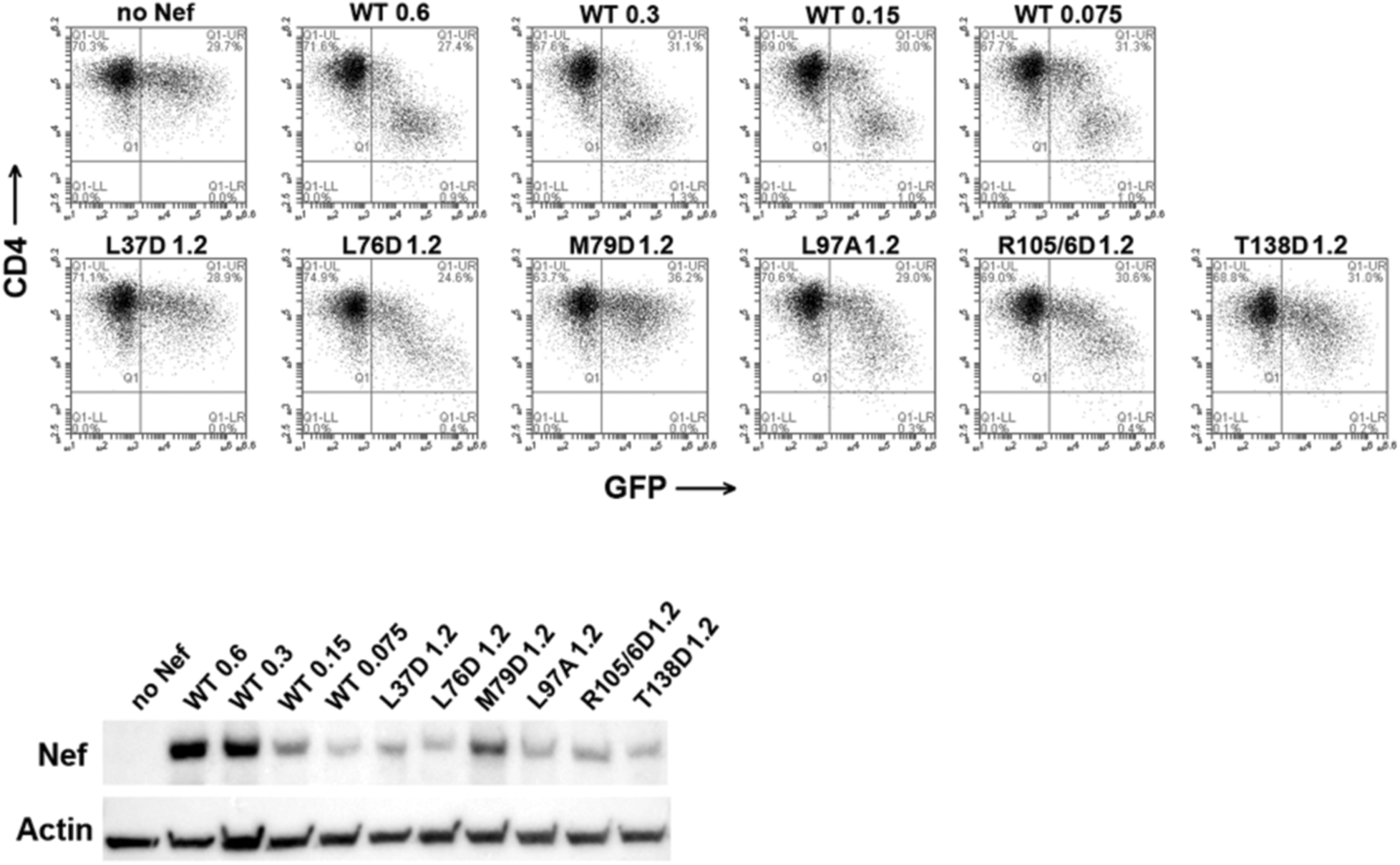
CD4 downregulation by the relatively poorly expressed Nef-mutants including a dose-response series for the expression of the wild type. HeLa cells that stably express CD4 were transfected to express GFP (a transfection marker) and Nef or the indicated Nef-mutants. Top: two color flow cytometry. Bottom: immunoblot for Nef and Actin. Numbers indicate the amount of Nef-expression plasmid in micrograms. Wild type (WT) Nef was expressed in a series of decreasing plasmid amounts to allow comparisons to the relatively poorly expressed Nef-mutants. The L37D and M79D substitutions are predicted by the structure to impair interaction with CD4, whereas the L76D substitution is not. The functional defects of the mutants L37D and M79D do not seem solely attributable to impaired expression, supporting the modeling of these residues as interacting with the cytoplasmic domain of CD4. L97A and R105/6D are mutants of the N-terminal-β-helix binding-site.

**Extended Data Fig. 2.**
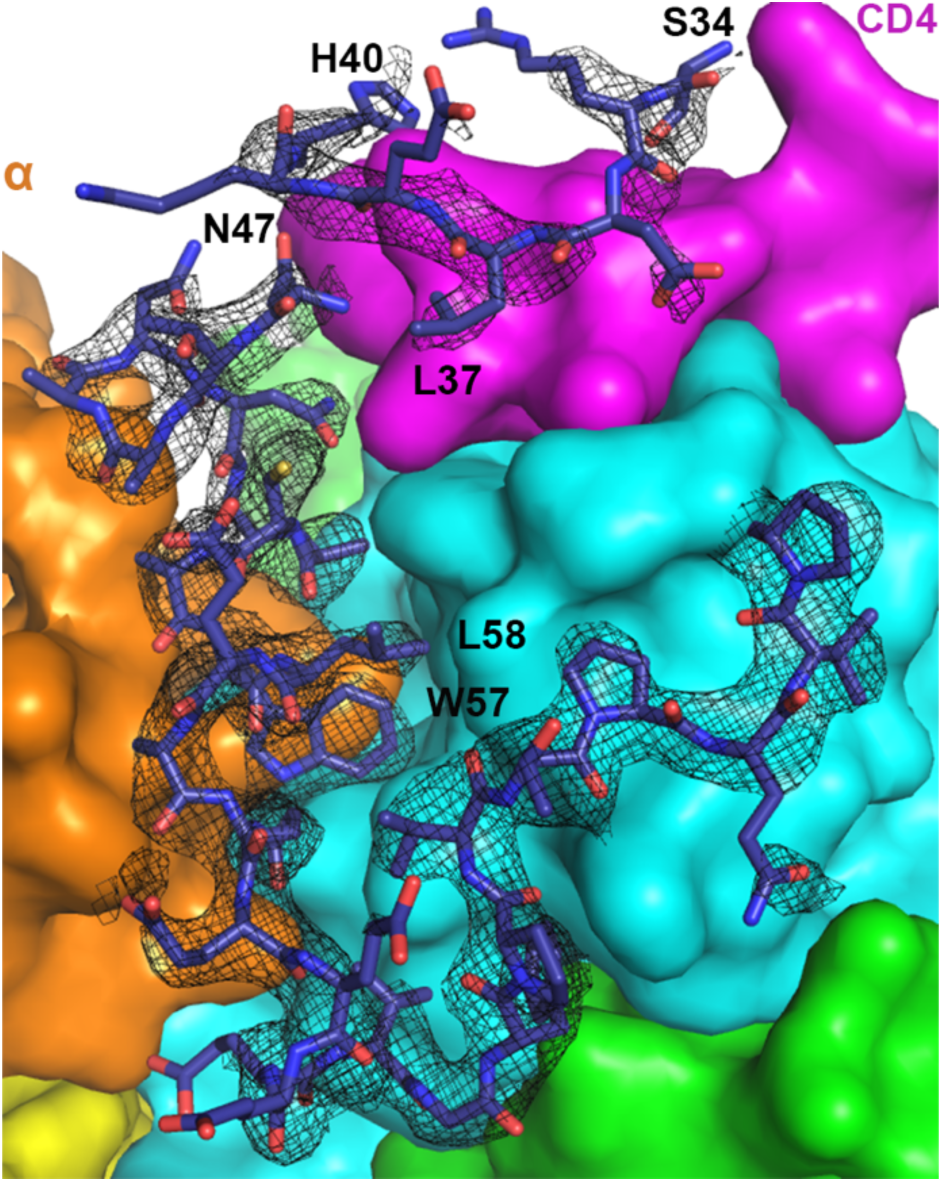
Electron density map for the N-terminal loop of Nef. 2Fo-Fc map (1σ level with B factor sharpened by −50Å^2^) for Nef residues 34-40 and 47-75 is shown as black mesh. Nef residues 41-46 could not be built due to the lack of density. Density for Nef 34-40 is less defined, although sidechain density for Leu37 is clear.

**Extended Data Fig. 3.**
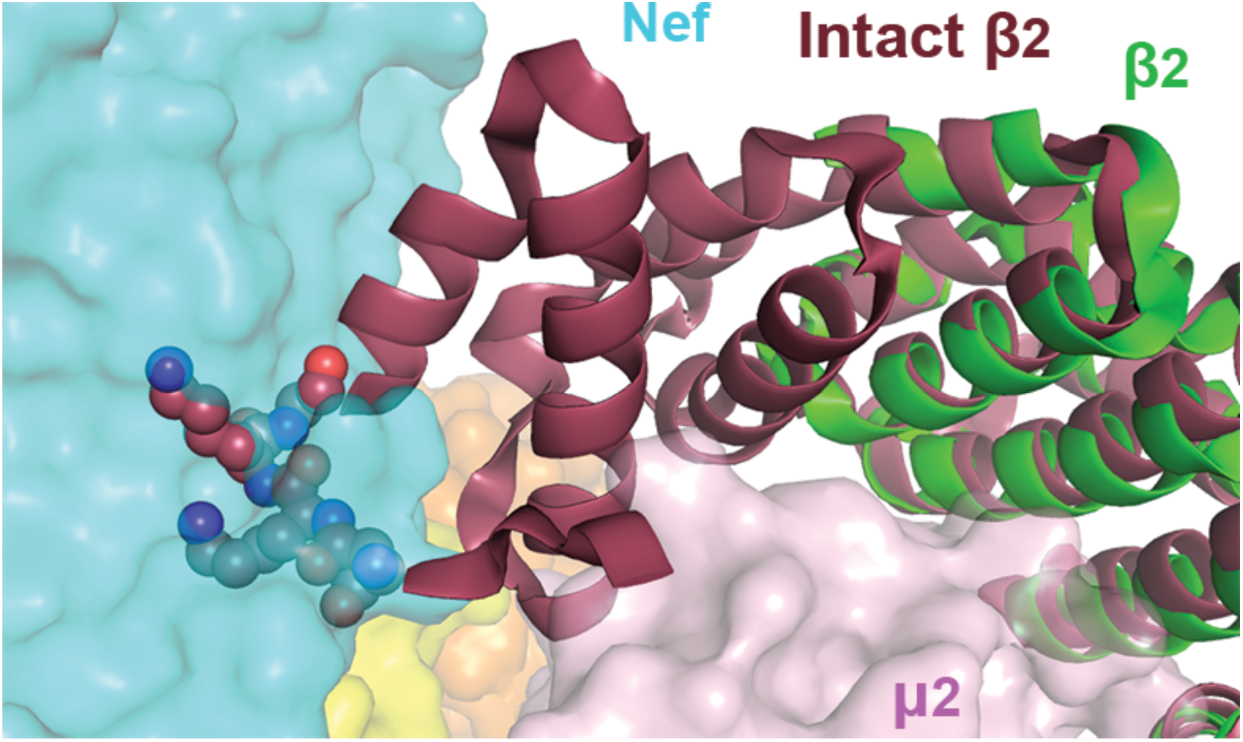
β2 subunit, if intact, would clash with the bound Nef. Overlay of the intact β2 subunit (dark red, 2XA7) with β2 in the current structure (green) indicates that clashing would take place between Nef and N-terminus of the intact β2, specifically residues Asn10, Lys11, Lys12, and Gly13 (spheres).

**Extended Data Fig. 4.**
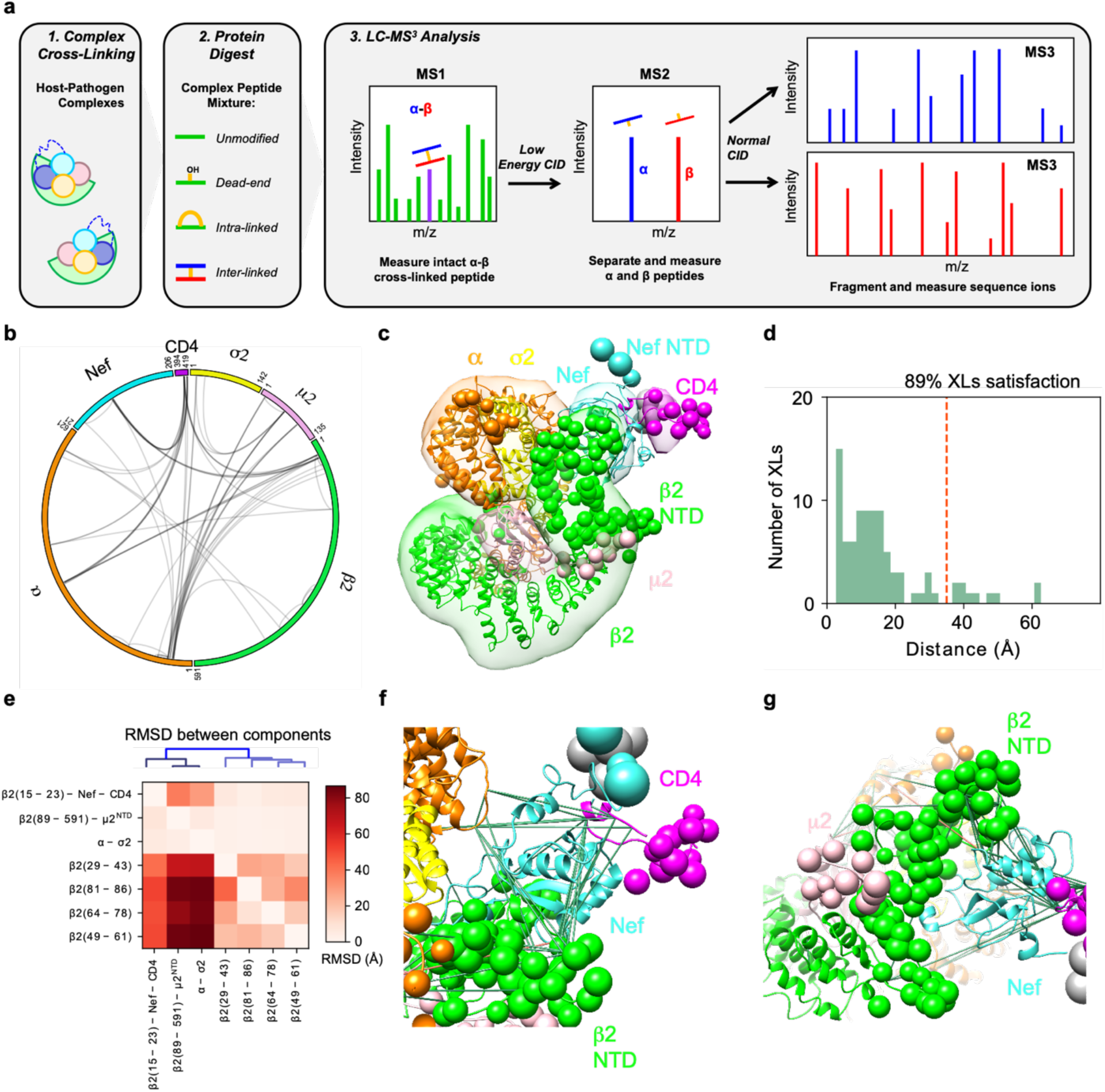
Crosslinking mass spectrometry and integrative structure modeling of the Nef-CD4_CD_-AP2^Δμ2-CTD^ complex. **a**, Overview of the DSSO XL-MS^3^ analysis method. **b**, CX-Circos linkage map of all Nef-CD4_CD_-AP2^Δμ2-CTD^ interlinks. **c**, Integrative structure of the Nef-CD4_CD_-AP2^Δμ2-CTD^ complex. The localization probability density of the ensemble of structures is shown with representative (centroid) structure from the ensemble embedded within it. Regions present in the crystal structure are shown as ribbons and segments not present in the crystal structure are shown as beads. **d**, Histogram showing the distribution of the cross-linked Cα–Cα distances in the integrative structure. The structural ensemble satisfies 89% of the XLs used to compute it. **e**, RMSD between rigid-bodies in the model ensemble. The vertical axis corresponds to the rigid body used as reference for superimposition and the horizontal axis are the rigid bodies for which the average RMSD was computed. **f**, Detail of crosslinks mapped to Nef. Satisfied and violated crosslinks shown in green and pink, respectively. **g**, Positioning of the unfolded β2 segment.

**Extended Data Fig. 5.**
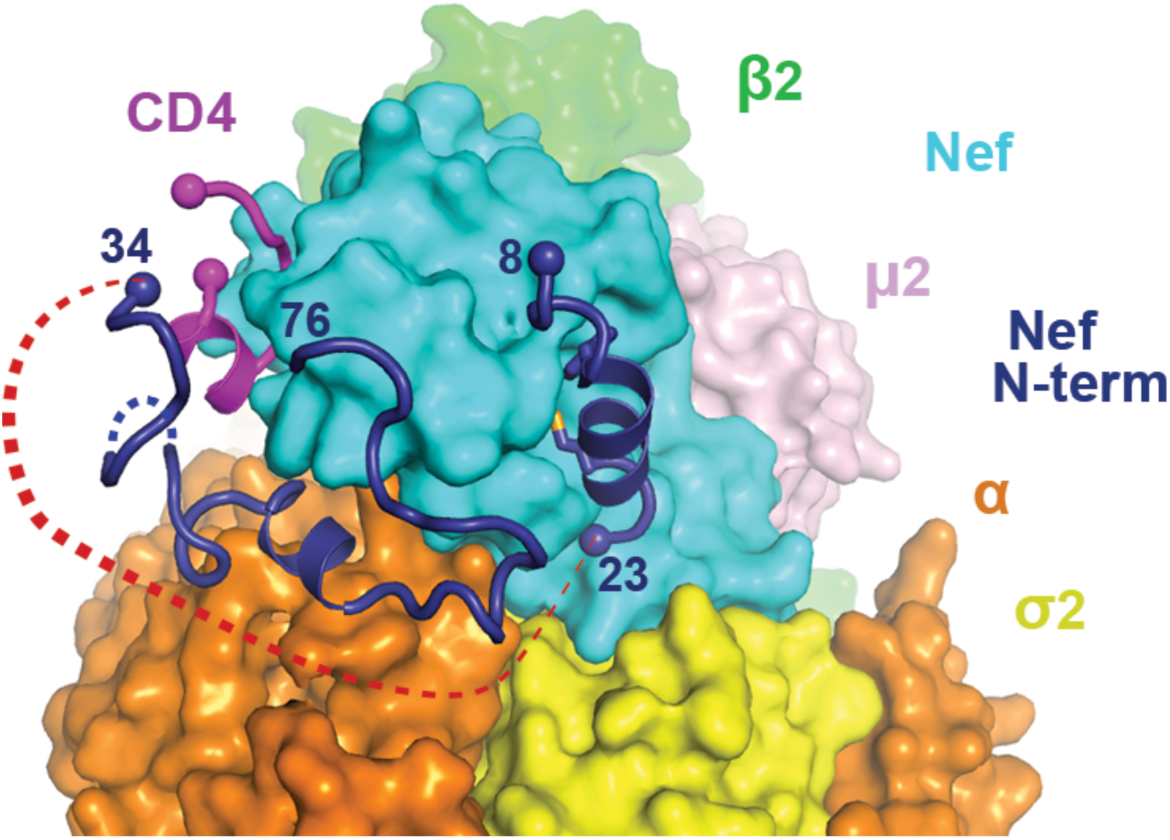
Binding of Nef N-terminal helix to the Nef core is incompatible with CD4 downregulation. N-terminal helix of Nef (8-23) is modeled into the current structure. Red dotted line represents the would-be distance between residues 23 and 34, which cannot be covered by ten residues (Nef 24-33).

**Extended Data Fig. 6.**
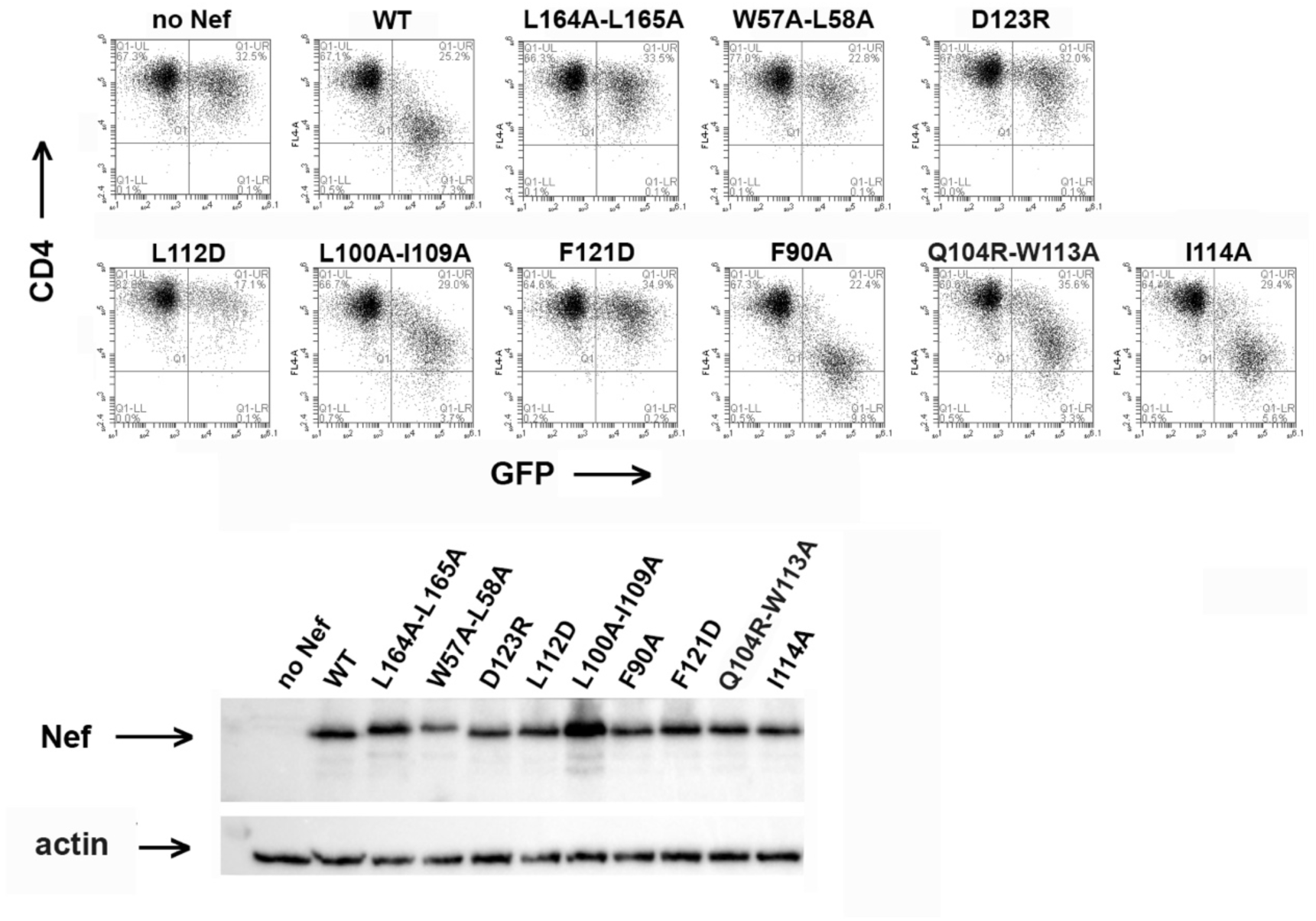
Mutations lining the helix-binding site on Nef minimally affect CD4 downregulation. HeLa cells that stably express CD4 were transfected to express GFP (a transfection marker) and Nef or the indicated Nef-mutants. Top: two color flow cytometry. Bottom: immunoblot for Nef and Actin. All the Nef-mutants of the N-terminal β-helix binding site (L100A-I109A, F90A, Q104R-W113A, and I114A) were well-expressed and were minimally if at all impaired for CD4-downregulation.

**Extended Data Fig. 7.**
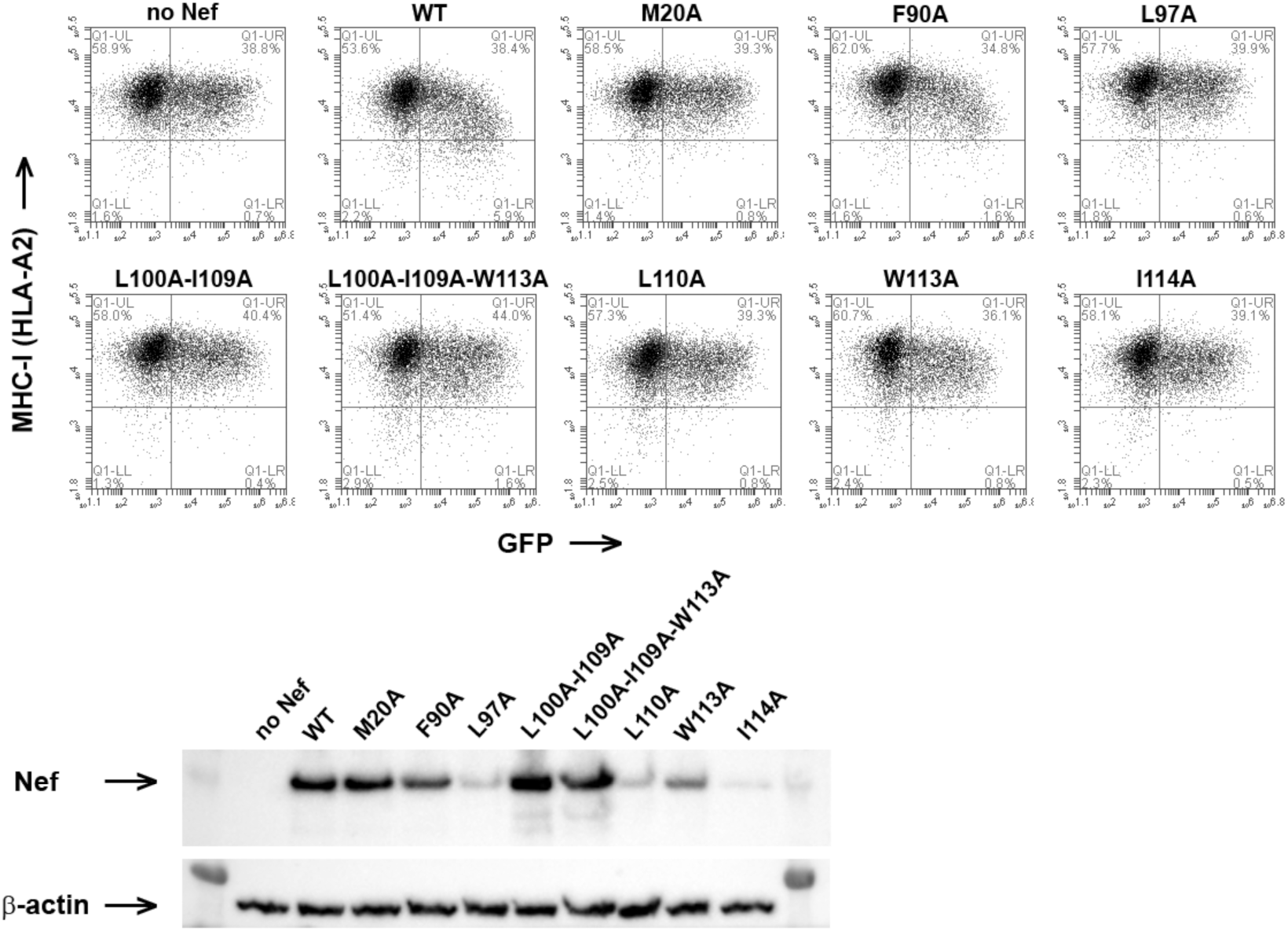
Mutations lining the helix-binding site on Nef affect class I MHC downregulation. HEK293 cells, which are naturally HLA-A2 positive, were transfected to express GFP (a transfection marker) and Nef or the indicated Nef-mutants. Top: two color flow cytometry. Bottom: immunoblot for Nef and Actin. The Nef-mutants of the N-terminal β-helix binding site with the exception of F90A were unable to downregulate class I MHC. Most of the mutants were poorly expressed, but L100A-I109A and L100A-I109A-W113A were well expressed yet functionally defective. M20A is a well-described mutant that is unable to downregulate class I MHC.

**Extended Data Fig. 8.**
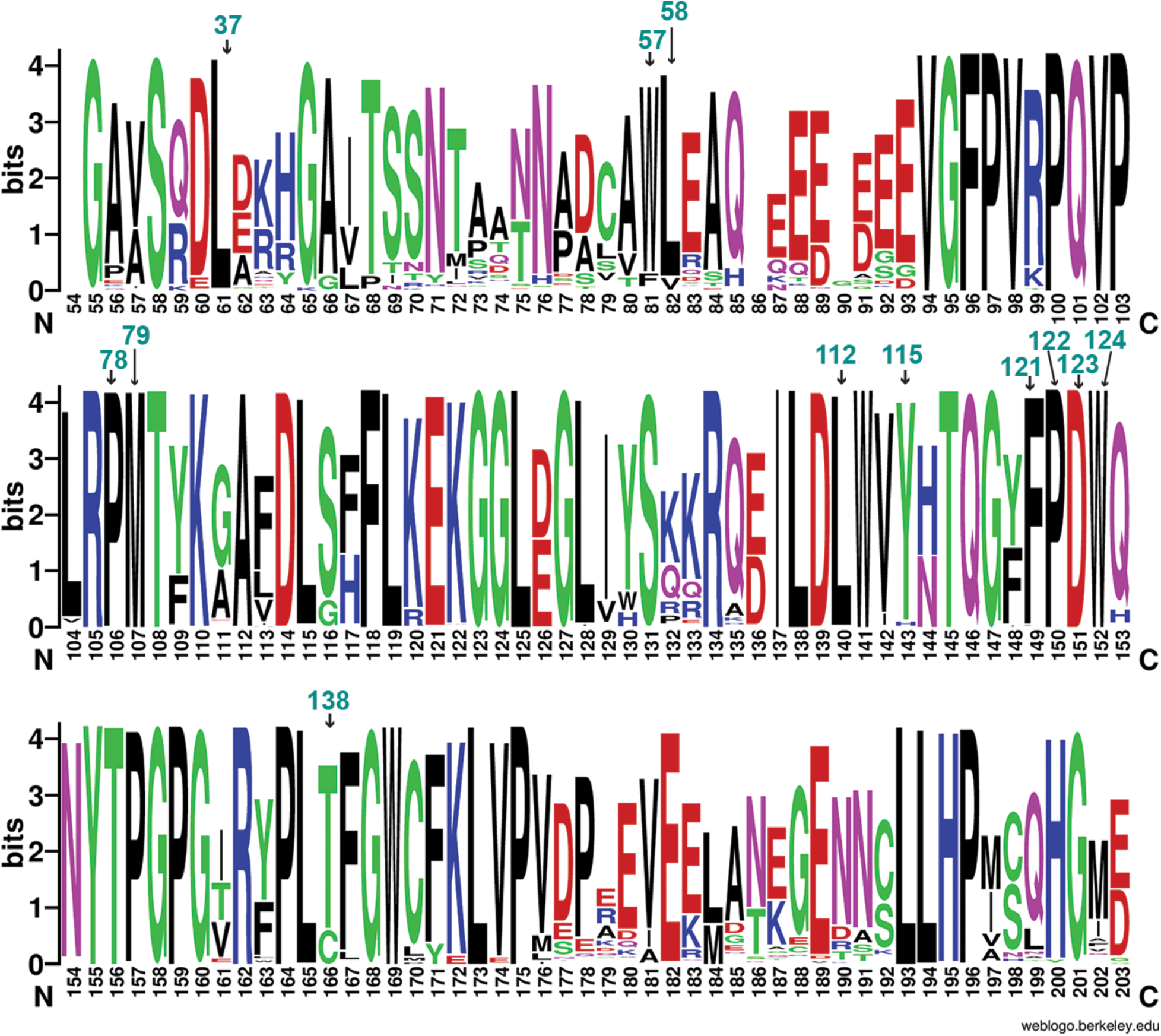
Nef residues important for CD4 downregulation are highly conserved. Nef sequences from HIV sequence compendium 2017 were analyzed though multiple sequence alignment (HIV sequence database, www.hiv.lanl.gov). Alignment was done in HXB2 convention (bottom) and residues important for CD4 downregulation are additionally labeled using the NL4.3 convention (top, cyan texts). The logo representation, with the height of each letter proportional to the observed frequency of the corresponding amino acid residue, was generated by WebLogo ^59^.

**Extended Data Table 1.** Crystallographic data collection and refinement statistics.

|  | Native |
| --- | --- |
| **Data collection** |  |
| Space group | P4<sub>1</sub> |
| Cell dimensions |  |
| *a*, *b*, *c* (Å) | 109.2, 109.2, 178.9 |
| Wavelength (Å) | 0.9793 |
| Resolution (Å) | 47.2 - 3.0 (3.11 - 3.0) |
| *R*<sub>merge</sub> | 0.287 |
| *I* / $\sigma I$ | 11.8 (1.4) |
| Completeness (%) | 100 (100) |
| Redundancy | 16.0 (15.9) |
| **Refinement** |  |
| Unique reflections | 41865 (4189) |
| *R*<sub>work</sub> / *R*<sub>free</sub> | 0.241/0.277 |
| No. atoms |  |
| Protein | 12261 |
| Water | 2 |
| *B*-factors |  |
| Protein | 118.30 |
| Water | 132.22 |
| R.m.s deviations |  |
| Bond lengths (Å) | 0.005 |
| Bond angles (°) | 0.88 |
| Ramachandran |  |
| Favored | 96.42% |
| Outliers | 0.13% |

\*Values in parenthesis are for the highest-resolution shell.

**Extended Data Table 2.**
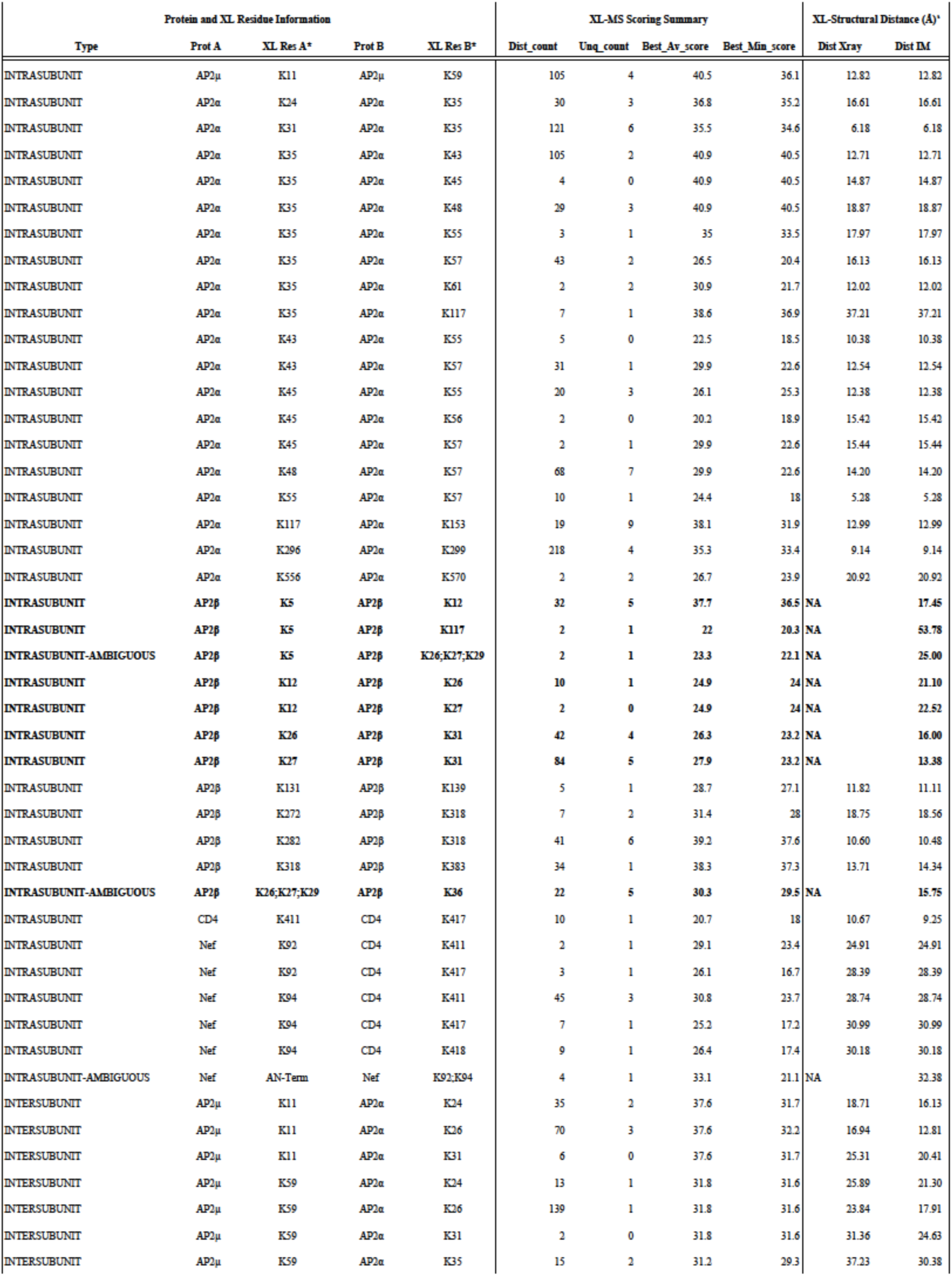
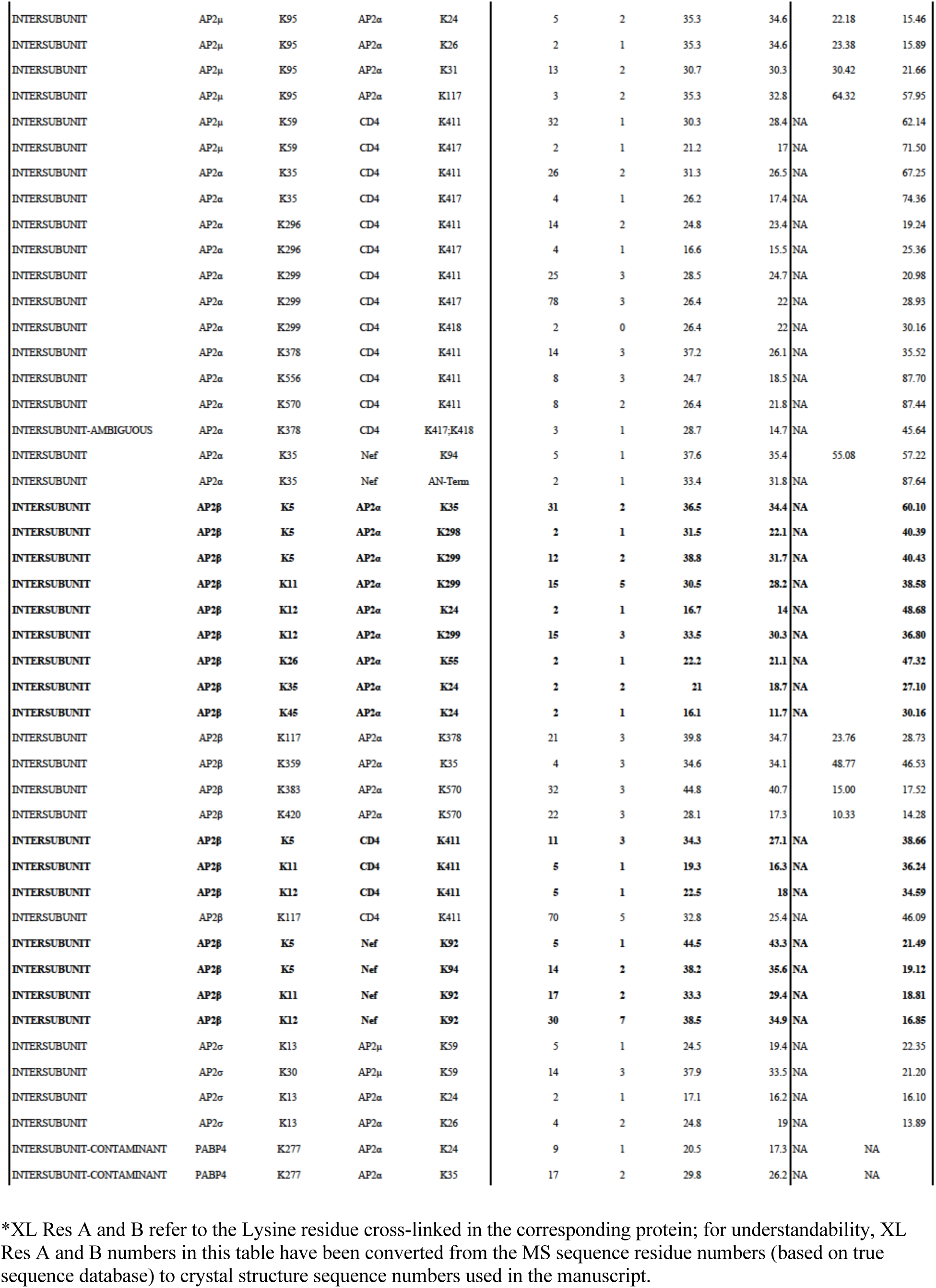
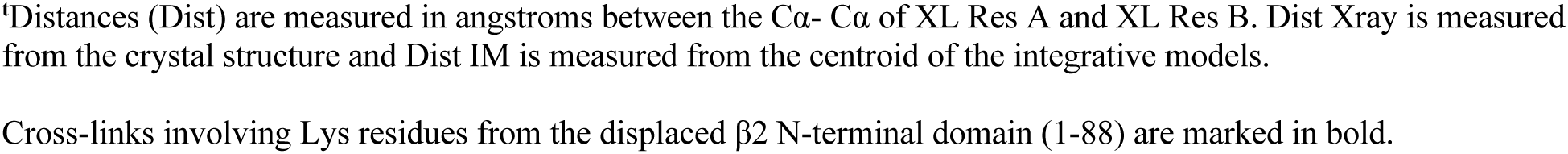
Intra- and inter-subunit DSSO inter-linked residues of Nef-CD4CD-AP2Δμ2-CTD.

| Protein and XL Residue Information |  |  |  |  | XL-MS Scoring Summary |  |  |  | XL-Structural Distance (Å)<sup>a</sup> |  |
| --- | --- | --- | --- | --- | --- | --- | --- | --- | --- | --- |
| Type | Prot A | XL Res A* | Prot B | XL Res B* | Dist_count | Unq_count | Best_Av_score | Best_Min_score | Dist Xray | Dist IM |
| INTRASUBUNIT | AP2μ | K11 | AP2μ | K59 | 105 | 4 | 40.5 | 36.1 | 12.82 | 12.82 |
| INTRASUBUNIT | AP2α | K24 | AP2α | K35 | 30 | 3 | 36.8 | 35.2 | 16.61 | 16.61 |
| INTRASUBUNIT | AP2α | K31 | AP2α | K35 | 121 | 6 | 35.5 | 34.6 | 6.18 | 6.18 |
| INTRASUBUNIT | AP2α | K35 | AP2α | K43 | 105 | 2 | 40.9 | 40.5 | 12.71 | 12.71 |
| INTRASUBUNIT | AP2α | K35 | AP2α | K45 | 4 | 0 | 40.9 | 40.5 | 14.87 | 14.87 |
| INTRASUBUNIT | AP2α | K35 | AP2α | K48 | 29 | 3 | 40.9 | 40.5 | 18.87 | 18.87 |
| INTRASUBUNIT | AP2α | K35 | AP2α | K55 | 3 | 1 | 35 | 33.5 | 17.97 | 17.97 |
| INTRASUBUNIT | AP2α | K35 | AP2α | K57 | 43 | 2 | 26.5 | 20.4 | 16.13 | 16.13 |
| INTRASUBUNIT | AP2α | K35 | AP2α | K61 | 2 | 2 | 30.9 | 21.7 | 12.02 | 12.02 |
| INTRASUBUNIT | AP2α | K35 | AP2α | K117 | 7 | 1 | 38.6 | 36.9 | 37.21 | 37.21 |
| INTRASUBUNIT | AP2α | K43 | AP2α | K55 | 5 | 0 | 22.5 | 18.5 | 10.38 | 10.38 |
| INTRASUBUNIT | AP2α | K43 | AP2α | K57 | 31 | 1 | 29.9 | 22.6 | 12.54 | 12.54 |
| INTRASUBUNIT | AP2α | K45 | AP2α | K55 | 20 | 3 | 26.1 | 25.3 | 12.38 | 12.38 |
| INTRASUBUNIT | AP2α | K45 | AP2α | K56 | 2 | 0 | 20.2 | 18.9 | 15.42 | 15.42 |
| INTRASUBUNIT | AP2α | K45 | AP2α | K57 | 2 | 1 | 29.9 | 22.6 | 15.44 | 15.44 |
| INTRASUBUNIT | AP2α | K48 | AP2α | K57 | 68 | 7 | 29.9 | 22.6 | 14.20 | 14.20 |
| INTRASUBUNIT | AP2α | K55 | AP2α | K57 | 10 | 1 | 24.4 | 18 | 5.28 | 5.28 |
| INTRASUBUNIT | AP2α | K117 | AP2α | K153 | 19 | 9 | 38.1 | 31.9 | 12.99 | 12.99 |
| INTRASUBUNIT | AP2α | K296 | AP2α | K299 | 218 | 4 | 35.3 | 33.4 | 9.14 | 9.14 |
| INTRASUBUNIT | AP2α | K556 | AP2α | K570 | 2 | 2 | 26.7 | 23.9 | 20.92 | 20.92 |
| INTRASUBUNIT | AP2β | K5 | AP2β | K12 | 32 | 5 | 37.7 | 36.5 | NA | 17.45 |
| INTRASUBUNIT | AP2β | K5 | AP2β | K117 | 2 | 1 | 22 | 20.3 | NA | 53.78 |
| INTRASUBUNIT-AMBIGUOUS | AP2β | K5 | AP2β | K26;K27;K29 | 2 | 1 | 23.3 | 22.1 | NA | 25.00 |
| INTRASUBUNIT | AP2β | K12 | AP2β | K26 | 10 | 1 | 24.9 | 24 | NA | 21.10 |
| INTRASUBUNIT | AP2β | K12 | AP2β | K27 | 2 | 0 | 24.9 | 24 | NA | 22.52 |
| INTRASUBUNIT | AP2β | K26 | AP2β | K31 | 42 | 4 | 26.3 | 23.2 | NA | 16.00 |
| INTRASUBUNIT | AP2β | K27 | AP2β | K31 | 84 | 5 | 27.9 | 23.2 | NA | 13.38 |
| INTRASUBUNIT | AP2β | K131 | AP2β | K139 | 5 | 1 | 28.7 | 27.1 | 11.82 | 11.11 |
| INTRASUBUNIT | AP2β | K272 | AP2β | K318 | 7 | 2 | 31.4 | 28 | 18.75 | 18.56 |
| INTRASUBUNIT | AP2β | K282 | AP2β | K318 | 41 | 6 | 39.2 | 37.6 | 10.60 | 10.48 |
| INTRASUBUNIT | AP2β | K318 | AP2β | K383 | 34 | 1 | 38.3 | 37.3 | 13.71 | 14.34 |
| INTRASUBUNIT-AMBIGUOUS | AP2β | K26;K27;K29 | AP2β | K36 | 22 | 5 | 30.3 | 29.5 | NA | 15.75 |
| INTRASUBUNIT | CD4 | K411 | CD4 | K417 | 10 | 1 | 20.7 | 18 | 10.67 | 9.25 |
| INTRASUBUNIT | Nef | K92 | CD4 | K411 | 2 | 1 | 29.1 | 23.4 | 24.91 | 24.91 |
| INTRASUBUNIT | Nef | K92 | CD4 | K417 | 3 | 1 | 26.1 | 16.7 | 28.39 | 28.39 |
| INTRASUBUNIT | Nef | K94 | CD4 | K411 | 45 | 3 | 30.8 | 23.7 | 28.74 | 28.74 |
| INTRASUBUNIT | Nef | K94 | CD4 | K417 | 7 | 1 | 25.2 | 17.2 | 30.99 | 30.99 |
| INTRASUBUNIT | Nef | K94 | CD4 | K418 | 9 | 1 | 26.4 | 17.4 | 30.18 | 30.18 |
| INTRASUBUNIT-AMBIGUOUS | Nef | AN-Term | Nef | K92;K94 | 4 | 1 | 33.1 | 21.1 | NA | 32.38 |
| INTERSUBUNIT | AP2μ | K11 | AP2α | K24 | 35 | 2 | 37.6 | 31.7 | 18.71 | 16.13 |
| INTERSUBUNIT | AP2μ | K11 | AP2α | K26 | 70 | 3 | 37.6 | 32.2 | 16.94 | 12.81 |
| INTERSUBUNIT | AP2μ | K11 | AP2α | K31 | 6 | 0 | 37.6 | 31.7 | 25.31 | 20.41 |
| INTERSUBUNIT | AP2μ | K59 | AP2α | K24 | 13 | 1 | 31.8 | 31.6 | 25.89 | 21.30 |
| INTERSUBUNIT | AP2μ | K59 | AP2α | K26 | 139 | 1 | 31.8 | 31.6 | 23.84 | 17.91 |
| INTERSUBUNIT | AP2μ | K59 | AP2α | K31 | 2 | 0 | 31.8 | 31.6 | 31.36 | 24.63 |
| INTERSUBUNIT | AP2μ | K59 | AP2α | K35 | 15 | 2 | 31.2 | 29.3 | 37.23 | 30.38 |
| INTERSUBUNIT | AP2μ | K95 | AP2α | K24 | 5 | 2 | 35.3 | 34.6 | 22.18 | 15.46 |
| INTERSUBUNIT | AP2μ | K95 | AP2α | K26 | 2 | 1 | 35.3 | 34.6 | 23.38 | 15.89 |
| INTERSUBUNIT | AP2μ | K95 | AP2α | K31 | 13 | 2 | 30.7 | 30.3 | 30.42 | 21.66 |
| INTERSUBUNIT | AP2μ | K95 | AP2α | K117 | 3 | 2 | 35.3 | 32.8 | 64.32 | 57.95 |
| INTERSUBUNIT | AP2μ | K59 | CD4 | K411 | 32 | 1 | 30.3 | 28.4 | NA | 62.14 |
| INTERSUBUNIT | AP2μ | K59 | CD4 | K417 | 2 | 1 | 21.2 | 17 | NA | 71.50 |
| INTERSUBUNIT | AP2α | K35 | CD4 | K411 | 26 | 2 | 31.3 | 26.5 | NA | 67.25 |
| INTERSUBUNIT | AP2α | K35 | CD4 | K417 | 4 | 1 | 26.2 | 17.4 | NA | 74.36 |
| INTERSUBUNIT | AP2α | K296 | CD4 | K411 | 14 | 2 | 24.8 | 23.4 | NA | 19.24 |
| INTERSUBUNIT | AP2α | K296 | CD4 | K417 | 4 | 1 | 16.6 | 15.5 | NA | 25.36 |
| INTERSUBUNIT | AP2α | K299 | CD4 | K411 | 25 | 3 | 28.5 | 24.7 | NA | 20.98 |
| INTERSUBUNIT | AP2α | K299 | CD4 | K417 | 78 | 3 | 26.4 | 22 | NA | 28.93 |
| INTERSUBUNIT | AP2α | K299 | CD4 | K418 | 2 | 0 | 26.4 | 22 | NA | 30.16 |
| INTERSUBUNIT | AP2α | K378 | CD4 | K411 | 14 | 3 | 37.2 | 26.1 | NA | 35.52 |
| INTERSUBUNIT | AP2α | K556 | CD4 | K411 | 8 | 3 | 24.7 | 18.5 | NA | 87.70 |
| INTERSUBUNIT | AP2α | K570 | CD4 | K411 | 8 | 2 | 26.4 | 21.8 | NA | 87.44 |
| INTERSUBUNIT-AMBIGUOUS | AP2α | K378 | CD4 | K417;K418 | 3 | 1 | 28.7 | 14.7 | NA | 45.64 |
| INTERSUBUNIT | AP2α | K35 | Nef | K94 | 5 | 1 | 37.6 | 35.4 | 55.08 | 57.22 |
| INTERSUBUNIT | AP2α | K35 | Nef | AN-Term | 2 | 1 | 33.4 | 31.8 | NA | 87.64 |
| INTERSUBUNIT | AP2β | K5 | AP2α | K35 | 31 | 2 | 36.5 | 34.4 | NA | 60.10 |
| INTERSUBUNIT | AP2β | K5 | AP2α | K298 | 2 | 1 | 31.5 | 22.1 | NA | 40.39 |
| INTERSUBUNIT | AP2β | K5 | AP2α | K299 | 12 | 2 | 38.8 | 31.7 | NA | 40.43 |
| INTERSUBUNIT | AP2β | K11 | AP2α | K299 | 15 | 5 | 30.5 | 28.2 | NA | 38.58 |
| INTERSUBUNIT | AP2β | K12 | AP2α | K24 | 2 | 1 | 16.7 | 14 | NA | 48.68 |
| INTERSUBUNIT | AP2β | K12 | AP2α | K299 | 15 | 3 | 33.5 | 30.3 | NA | 36.80 |
| INTERSUBUNIT | AP2β | K26 | AP2α | K55 | 2 | 1 | 22.2 | 21.1 | NA | 47.32 |
| INTERSUBUNIT | AP2β | K35 | AP2α | K24 | 2 | 2 | 21 | 18.7 | NA | 27.10 |
| INTERSUBUNIT | AP2β | K45 | AP2α | K24 | 2 | 1 | 16.1 | 11.7 | NA | 30.16 |
| INTERSUBUNIT | AP2β | K117 | AP2α | K378 | 21 | 3 | 39.8 | 34.7 | 23.76 | 28.73 |
| INTERSUBUNIT | AP2β | K359 | AP2α | K35 | 4 | 3 | 34.6 | 34.1 | 48.77 | 46.53 |
| INTERSUBUNIT | AP2β | K383 | AP2α | K570 | 32 | 3 | 44.8 | 40.7 | 15.00 | 17.52 |
| INTERSUBUNIT | AP2β | K420 | AP2α | K570 | 22 | 3 | 28.1 | 17.3 | 10.33 | 14.28 |
| INTERSUBUNIT | AP2β | K5 | CD4 | K411 | 11 | 3 | 34.3 | 27.1 | NA | 38.66 |
| INTERSUBUNIT | AP2β | K11 | CD4 | K411 | 5 | 1 | 19.3 | 16.3 | NA | 36.24 |
| INTERSUBUNIT | AP2β | K12 | CD4 | K411 | 5 | 1 | 22.5 | 18 | NA | 34.59 |
| INTERSUBUNIT | AP2β | K117 | CD4 | K411 | 70 | 5 | 32.8 | 25.4 | NA | 46.09 |
| INTERSUBUNIT | AP2β | K5 | Nef | K92 | 5 | 1 | 44.5 | 43.3 | NA | 21.49 |
| INTERSUBUNIT | AP2β | K5 | Nef | K94 | 14 | 2 | 38.2 | 35.6 | NA | 19.12 |
| INTERSUBUNIT | AP2β | K11 | Nef | K92 | 17 | 2 | 33.3 | 29.4 | NA | 18.81 |
| INTERSUBUNIT | AP2β | K12 | Nef | K92 | 30 | 7 | 38.5 | 34.9 | NA | 16.85 |
| INTERSUBUNIT | AP2σ | K13 | AP2μ | K59 | 5 | 1 | 24.5 | 19.4 | NA | 22.35 |
| INTERSUBUNIT | AP2σ | K30 | AP2μ | K59 | 14 | 3 | 37.9 | 33.5 | NA | 21.20 |
| INTERSUBUNIT | AP2σ | K13 | AP2α | K24 | 2 | 1 | 17.1 | 16.2 | NA | 16.10 |
| INTERSUBUNIT | AP2σ | K13 | AP2α | K26 | 4 | 2 | 24.8 | 19 | NA | 13.89 |
| INTERSUBUNIT-CONTAMINANT | PABP4 | K277 | AP2α | K24 | 9 | 1 | 20.5 | 17.3 | NA | NA |
| INTERSUBUNIT-CONTAMINANT | PABP4 | K277 | AP2α | K35 | 17 | 2 | 29.8 | 26.2 | NA | NA |

\*XL Res A and B refer to the Lysine residue cross-linked in the corresponding protein; for understandability, XL Res A and B numbers in this table have been converted from the MS sequence residue numbers (based on true sequence database) to crystal structure sequence numbers used in the manuscript.
<sup>†</sup>Distances (Dist) are measured in angstroms between the Cα - Cα of XL Res A and XL Res B. Dist Xray is measured from the crystal structure and Dist IM is measured from the centroid of the integrative models.
- 5 Cross-links involving Lys residues from the displaced β2 N-terminal domain (1-88) are marked in bold.

